# A Whole-Genome and Ancient DNA Perspective on the Drivers of Genetic Diversity and Structure in Palearctic True Lemmings

**DOI:** 10.64898/2026.06.15.731284

**Authors:** Ivan Dvoyashov, Tatyana Petrova, Valentina Panitsina, Semyon Bodrov, Natalia Serdyuk, Albert Protopopov, Aisen Klimovskiy, Mikhail Tiunov, Alexey Lopatin, Leonid Lavrenchenko, Natalia Abramson

**Affiliations:** A.N. Severtsov Institute of Ecology and Evolution, Russian Academy of Sciences, 119071 Moscow, Russia; Zoological Institute of the Russian Academy of Sciences, 199034 St. Petersburg, Russia; Borissiak Paleontological Institute of the Russian Academy of Sciences, 117647 Moscow, Russia; Academy of Sciences of the Republic of Sakha (Yakutia), 677007 Yakutsk, Russia; Federal Scientific Center of the East Asia Terrestrial Biodiversity of the Far Eastern Branch of the Russian Academy of Sciences, 690022 Vladivostok, Russia

**Keywords:** *Lemmus*, isolation-by-distance, phylogenomics, Arctic biodiversity, postglacial colonisation, Last Glacial Maximum

## Abstract

True lemmings (genus *Lemmus*) underwent substantial range shifts during the Late Pleistocene and the Pleistocene–Holocene transition, but the impact of these events on present-day diversity remains poorly understood. Here, we used whole-genome sequencing data from modern and ancient samples across the Palearctic range to address this knowledge gap. Reconstruction of autosomal phylogeny revealed that Palearctic true lemmings exhibit relatively shallow genetic structure, contrasting with the deep divergence inferred from mitochondrial genomes. Genetic variation largely follows an isolation-by-distance pattern, and no elevated nuclear divergence was detected between the major mitochondrial lineages. Window-based phylogenetic analyses identified several peripheral populations with high concordance factors, including Norway and Amur lemmings. The high degree of phylogenetic concordance along the genome in these populations is likely a consequence of postglacial bottlenecks and isolation, as indicated by reduced heterozygosity and the presence of runs of homozygosity in them. Overall, our results indicate that the modern genomic structure of Palearctic lemmings was shaped primarily by range fragmentation and population isolation following the broad distribution of the genus during the Last Glacial Maximum. Thus, the current genetic structure appears to represent only a fraction of the Late Pleistocene true lemming diversity. This is illustrated by a genetically distinct ancient specimen (∼40 ka BP) from the Indigirka River basin that does not cluster with any modern lineage. From a taxonomic perspective, these findings do not support strong species-level differentiation among the major Palearctic lineages and highlight the discrepancy between mitochondrial and nuclear patterns of diversity within the genus.

## 1 Introduction

True lemmings (genus *Lemmus* Link, 1795) today are a crucial component of Arctic ecosystems, together with collared lemmings forming a main prey base for Arctic predators such as snowy owls, Arctic foxes, stoats, and others (Legagneux et al., 2012). During the Pleistocene–Holocene climatic transition, lemmings underwent substantial range shifts that led to a significant reduction of genetic diversity in European populations (Lagerholm et al., 2014). Similar evolutionary patterns have also been described for other Arctic species, including the collared lemming *Dicrostonyx torquatus* (Pallas, 1778) and the narrow-headed vole *Lasiopodomys gregalis* (Pallas, 1779), leading to the formulation of the model of genetic extinction and diminution as a general response of Arctic species in Europe to climatic oscillations (Stojak and Jędrzejewska, 2022). However, considering the circumpolar distribution of lemmings, this model may not adequately reflect the evolutionary history of the genus across its entire range. Given that cold-adapted taxa are strongly affected by current global warming, knowledge of past biogeographic dynamics is essential for understanding their responses and for conserving modern biodiversity and habitats.

Knowledge of the distribution of the genus during the Last Glacial Maximum (LGM) and post-LGM period is most detailed for Europe, although some information is also available for Asia. Paleontological evidence indicates that during the LGM true lemmings occupied much of continental Europe, from the margin of the Scandinavian Ice Sheet in the north to approximately 48°N in the south (Markova et al., 2019). There is also a hypothesis of their persistence in Scandinavian refugia (Fedorov and Stenseth, 2001). Numerous true lemming remains dated to the LGM have been discovered in the Ural Mountains, including both the southern and central parts of the mountain range (Markova et al., 2019). In the Asian part of the continent, available evidence likewise suggests a much broader former distribution, extending along the southern margin of the West Siberian Lowland, the Upper Ob region, and the Kuznetsk–Salair province of southern Siberia (Markova et al., 1995). In the east, the range of true lemmings extended as far as the Sikhote-Alin Mountain Range (Tiunov and Panasenko, 2011).

Although still abundant in Europe during the Early Late Glacial (18–14.7 ka), true lemmings began to disappear during the Bølling–Allerød interstadial, while simultaneously colonizing northern territories released from beneath the Scandinavian Ice Sheet (Markova et al., 2019). Today, in the European part of the continent, they persist only in Scandinavia, on the Kola Peninsula, and in lowland and high-altitude tundra extending from the Kanin Peninsula to the Ural Mountains. A similar range contraction occurred in Asia, where true lemmings are now abundant mainly in the high-latitude lowland tundra from the Yamal Peninsula to the Chukotka Peninsula. Nevertheless, small isolated populations recorded during the 20th century, much farther south — in the Verkhoyansk Range, eastern Transbaikalia, the Amur region, and Kamchatka Peninsula — likely represent remnants of this formerly extensive distribution.

The complex history of range expansion, contraction, fragmentation, and secondary contact during the Late Pleistocene and Holocene played a major role in shaping the present-day genetic structure of true lemmings. Nevertheless, many aspects of the evolutionary relationships and taxonomic structure within the genus remain unresolved.

Based on mitochondrial cytochrome *b* (mt *cytb*) data, previous studies (Abramson and Petrova, 2018; Abramson et al., 2008; Fedorov et al., 1999) identified several deeply divergent monophyletic groups within the genus *Lemmus*. First, all true lemmings split into two major mt branches. The Nearctic branch is represented by *L. trimucronatus* (Richardson, 1825). Within the Palearctic branch, mt lineages broadly corresponding to *Lemmus amurensis* Vinogradov, 1924 and *L. lemmus* (Linnaeus, 1758) were identified. Also the paraphyly of *L. sibiricus* (Kerr, 1792) was revealed. The western lineage of *L. sibiricus* (sister to *L. lemmus*) occupies territories west of the Lena River, whereas the eastern lineage (sister to *L. amurensis*) occurs east of the Lena River delta and extends to the Kolyma River on the mainland; it is also present on the New Siberian Islands, Wrangel Island, and in an isolated locality on the eastern side of the Kamchatka Peninsula.

Additional information on the phylogeography and population history of true lemmings comes from time-calibrated mt phylogeny (Lord et al., 2025; Panitsina et al., 2026). Based on these analyses, the divergence time of the Norway lemming mt lineage was estimated at 20–18 ka, corresponding to the deglaciation of Scandinavia and the Kola Peninsula. The divergence between the eastern and western lineages of *L. sibiricus* was estimated at approximately 92 ka BP. The Amur lemming lineage is sister to the eastern lineage of *L. sibiricus* and probably diverged during the Late Pleistocene, predating the LGM.

Mitochondrial results support the species-level status of *L. amurensis* and *L. lemmus* and additionally raise the question of whether the western and eastern lineages of *L. sibiricus* should also be assigned separate species status. However, mitochondrial DNA (mtDNA) variation may retain signals of ancient polymorphism and therefore reflect deeper evolutionary processes rather than the more recent demographic history responsible for present-day population structure (Toews and Brelsford, 2012). At the same time, classical karyological (Gileva et al., 1984) and hybridization studies (Pokrovski et al., 1984) found no evidence of reproductive barriers within the Palearctic branch of the genus *Lemmus*, that is, among all Palearctic representatives except *L. trimucronatus*. This evidence led to an alternative interpretation of species boundaries within the genus, according to which only two species are recognized: *L. lemmus*, representing the Palearctic branch, and *L. trimucronatus*, representing the Nearctic branch (Kryštufek and Shenbrot, 2022). The only whole-genome study published to date (Lord et al., 2025) recovered the nuclear monophyly of *L. lemmus*. However, the sampling in that study was limited, and most of the Palearctic range of the genus was not represented.

In this study, we aimed to investigate the evolutionary relationships of Palearctic true lemmings and identify the biogeographic factors that shaped it based on the whole-genome sequencing (WGS) data of modern and ancient Late Pleistocene and Holocene samples.

## 2 Materials and Methods

### 2.1 Sampling

Samples were obtained from the tissue collection of the Evolutionary Genomics and Paleogenomics Laboratory, Zoological Institute of the Russian Academy of Sciences, ZIN RAS (modern, fresh, ethanol-fixed tissues), from the voucher collection of the Laboratory of Theriology, ZIN RAS (museum skin specimens), the Borissiak Paleontological Institute of the Russian Academy of Sciences (Indigirka3 ancient sample; Lopatin et al., 2019), the Federal Scientific Center of East Asia Terrestrial Biodiversity of the Far Eastern Branch of the Russian Academy of Sciences (Sikhote ancient sample), and from the Academy of Sciences of the Republic of Sakha (Yakutia) (Indigirka1 and Indigirka2 ancient samples, but see Results section 3.1).

Our dataset spans the entire contemporary geographic range of the genus *Lemmus* and includes several highly isolated populations, such as *L. amurensis* and lemmings from a volcanic caldera Uzon (Kamchatka Peninsula). We sequenced 33 *Lemmus* genomes and one *D. torquatus* genome (as an outgroup), with variable coverage reflecting specimen quality. We additionally incorporated data from Lord et al. (2025), including nine modern and two ancient samples.

### 2.2 Data Generation

Sequencing procedures were described in detail in our previous studies on true lemmings mt phylogeny (Panitsina et al., 2026). Briefly, DNA from modern and museum samples was extracted using a phenol–chloroform protocol (Barnett and Larson, 2012; Green and Sambrook, 2017). Ancient DNA was extracted using a silica-based method (Rohland and Hofreiter, 2007) with modifications from Panitsina et al. (2023).

The average library length across samples was ∼250 bp. Libraries were sequenced using DNBSEQ-G400 (paired-end, 2 × 150 bp) (MGI Tech Co., Shenzhen, China). Raw reads are deposited in the NCBI Sequence Read Archive under BioProject PRJNA590630.

### 2.3 DNA Quality Assessment, Aligning and Calling

Read quality and adapter contamination were assessed with FastQC v.0.11.9 (Andrews, 2010) and trimmed using BBDuk (https://sourceforge.net/projects/bbmap/). Contamination was evaluated using a custom Kraken2 (Wood et al., 2019) database comprising bacterial and fungal genomes, the human genome, and genomes of *D. torquatus*, *L. lemmus*, and *Microtus ochrogaster* (Wagner, 1842). For contaminated samples, only Arvicolinae reads were retained using extract_kraken_reads.py.

Reads were aligned to the *L. lemmus* reference genome (GCA_964027135.1), combined with the mt genome of the same specimen, using BWA-MEM v.0.7.17 (Li, 2013). Resulting BAM files were filtered with SAMtools v.1.18 (-F 2828 –f 2) (Li et al., 2009), and duplicates were marked. Alignment quality was assessed with Qualimap v.2.3 (García-Alcalde et al., 2012), and postmortem damage (PMD) patterns in ancient samples were evaluated using DamageProfiler v.1.1 (Neukamm et al., 2021).

Variant calling was performed with BCFtools v.1.21 (Li, 2011), with ploidy specified for autosomes, X and Y chromosomes (defined following Feinauer et al., 2026), and the mt genome. All sites, including monomorphic ones, were called, after which indels were excluded. During variant calling, only reads with MAPQ > 30 were used. The group-samples option was disabled so that each individual was treated independently, following the rationale of De Jong et al. (2023). In the resulting autosomal VCF, we retained only autosomal scaffolds (scaffolds 1–25, excluding scaffold 3 identified as X chromosome) and filtered out positions with extreme total depth (<0.5× or >2.3× the median depth).

### 2.4 Phylogenetic analyses

For autosomal phylogeny, the VCF was filtered using bcftools view (F_MISSING ≤ 0.2, MAC ≥ 3), retaining only biallelic single nucleotide polymorphisms (SNPs). Genome-wide pairwise Manhattan distances were computed using a custom R script, and a BioNJ species tree was constructed using the ape package (Paradis and Schliep, 2019). This approach allows correct estimation of genetic distance for ancient samples with high missing rate, as distance calculation uses only sites, genotyped in both samples of the comparison pair. In addition, distance-based phylogenetic reconstruction is appropriate for this dataset because the studied populations represent shallow evolutionary divergences, for which multiple and reverse substitutions are expected to be rare, while genome-wide patterns of allele sharing shaped by recent demographic history are likely to represent the major source of genetic variation.

To assess clade genomic concordance, phylogenies were reconstructed for non-overlapping 1 Mb windows across the genome. This relatively large window size was chosen to mitigate the effects of low coverage and missing data in some specimens. Clade concordance factors were quantified using prop.clades from the phangorn package (Schliep, 2011) and mapped onto the species tree.

A phylogenetic network was constructed using consensusNet command (prob = 0.25). Clade concordance analyses account for discordant phylogenetic signals in an autosome dataset, therefore we can estimate “the proportion of the genome for which a given clade is true” (Baum, 2007).

In Panitsina et al. (2026), mt genomes were analysed for the same samples, excluding Yamal1, which was sequenced later. For this sample, the mt genome was obtained as a consensus sequence from the BAM file using samtools consensus. It was added to the assembled mt genomes from Panitsina et al., and the sequences were aligned with MAFFT v.7.526 (Katoh et al., 2019). Maximum Likelihood phylogenetic reconstruction was performed with RAxML-NG v.1.2.2 (Stamatakis, 2006).

### 2.5 Genetic Structure and Gene Flow

From the VCF obtained during phylogenetic analyses samples of *L. trimucronatus* and outgroups were excluded. The autosomal VCF was filtered (F_MISSING ≤ 0.1, MAC ≥ 3) and then Principal Component Analysis (PCA) was performed using the SNPRelate package (Zheng et al., 2012). We did not perform linkage disequilibrium (LD) pruning in order to retain the maximum number of genotyped SNPs for our samples. Given the low coverage of the ancient samples, the application of strict missingness filters per SNP resulted in a sparse SNP distribution, thereby reducing the number of sites in LD.

To assess gene flow between populations, we used two methods. First, ancestry coefficients were estimated using the LEA package (Frichot and François, 2015). For this, we used the same VCF as for PCA. The optimal number of clusters (K = 2–6) was determined using the elbow method based on cross-entropy values (20 runs per K), with the optimal K defined as the point at which the curve reached a plateau.

Second, we calculated ABBA–BABA statistics (Durand et al., 2011) using Dsuite v.0.5 (Malinsky et al., 2021) on the VCF obtained during phylogenetic analyses (section 2.4). This classical site-based test allows the inclusion of low-coverage samples, although it lacks some of the inferential power of locus-based tests (Hibbins and Hahn, 2022). Two analyses were performed: one with *Dicrostonyx* as the outgroup and one with *L. trimucronatus*. The use of *Dicrostonyx* as an outgroup allowed us to evaluate potential gene flow between the Nearctic and Palearctic branches of the genus, whereas the use of *L. trimucronatus* as an outgroup provided higher resolution for estimating gene flow within the Palearctic branch. Statistical significance of D-statistics was assessed using the block-jackknife procedure implemented in Dsuite, which accounts for linkage disequilibrium by resampling genomic blocks and provides standard errors and Z-scores for each test. Based on D-test results, Fbranch statistics were calculated for autosomal species tree and visualized using dtools.py.

### 2.6 Runs of Homozygosity and Heterozygosity Estimation

Due to heterogeneous sequencing depth across samples, Runs of homozygosity (ROH) analyses were restricted to individuals with mean coverage >5x. To minimize false homozygous calls caused by allele dropout at low coverage, all genotypes with depth <4 were masked and treated as missing. This strategy converts low-confidence genotype calls into missing data, allowing the use of stringent ROH detection parameters while accommodating uneven coverage across samples. Specifically, using PLINK v1.9.0 (Purcell et al., 2007) we applied strict limits on heterozygous calls within sliding windows (--homozyg-window-snp 1000, –-homozyg-window-het 1), while allowing for missing genotypes (--homozyg-window-missing 450), thereby reducing false ROH inflation without fragmenting ROH due to stochastic dropout.

Per-sample heterozygosity was estimated from the autosomal VCF described in Section 2.3 as the proportion of heterozygous genotypes among all called genotypes, including monomorphic sites. This analysis was performed only for samples with mean depth coverage above 3× and included only sites with individual coverage above 2×, following the threshold proposed by de Jong et al. (2023), using a custom awk script.

### 2.7 Isolation by Distance

Pairwise geographic distances between samples were calculated as least-cost path distances by land using the gdistance package. To account for potential dispersal during periods of low sea level (e.g., the LGM), islands and Alaska were connected to the mainland via shortest paths in the land raster. This approach reflects a model of historical connectivity, under which present-day island populations are not treated as completely isolated but as having been linked via land bridges during glacial periods.

Geographic distances were compared with pairwise genetic distances estimated during the autosomal analysis. Ancient samples were excluded from this dataset. Correlation was assessed using a Mantel test with the VEGAN R package (Dixon, 2003). Additionally, residuals from linear regression between geographic and genetic distances were plotted against mean heterozygosity for each sample pair to evaluate the effect of reduced genetic variation on genetic distance. In the last case, only samples with estimated heterozygosity were used (see section 2.6).

### 2.8. Demographic Modelling for *Lemmus lemmus*

A demographic model requires the specification of multiple parameters, including the time intervals for changes in population size, effective population sizes themselves, the timing of population divergence (in two-population models), and migration rates. Here, we implemented GADMA (Noskova et al., 2020) to optimize these parameters for models based on the site frequency spectrum (SFS). GADMA provides global parameter optimization using a genetic algorithm implemented for several demographic modelling engines, of which we selected moments (Jouganous et al., 2017).

Demographic modelling requires reliable population assignment and sufficient sample sizes. Therefore, analyses were restricted to two populations of *L. lemmus* with adequate representation: Scandinavia (n = 3) and the Kola Peninsula (n = 7). Genotypes with sequencing depth <5× were masked, and sites containing missing data were excluded. The total length of the resulting VCF, including both monomorphic and biallelic sites, was used as the effective sequence length for demographic inference. The mutation rate was set to 8.7 × 10□□ substitutions/site/generation, following Wang et al. (2023), and generation time was set to 0.5 years.

Folded site frequency spectra (SFS) were generated using easySFS (https://github.com/isaacovercast/easySFS) without projection, as the chosen VCF filtering procedure did not leave any missing genotypes. To assess parameter uncertainty and account for linkage among SNPs, the VCF was block-bootstrapped using custom scripts to generate 100 replicate datasets, from which replicate SFSs were constructed. These replicate spectra were used as input for demographic inference in GADMA.

GADMA allows specification of the maximum number of demographic epochs occurring before and after population splitting in two-population models. We allowed the GADMA algorithm to model a maximum of two epochs before population splitting and one epoch after splitting, as a larger number of epochs could potentially lead to overfitting. GADMA was run with 20 independent replicates. The resulting models were evaluated based on log-likelihood values and the composite likelihood Akaike information criterion (CLAIC), which is appropriate for linked SNP data.

## 3 Results

### 3.1 Genetic Structure of the True Lemmings

Our final dataset comprised 44 samples from 32 localities of true lemmings (Fig. 1A; Table S1). Resulting coverage of our samples after all filtering steps was dependent on the degree of tissue preservation (Table S2). The final VCF used to construct genome-wide autosomal phylogeny and window-based phylogeny contained 137 M SNPs. The thinned dataset used to construct PCA, structure-like analyses implemented in LEA contained 2.5 M SNPs. Some general statistics about quality of the reads, alignment and contamination are present in Supplementary Table S2.

**Figure 1.**
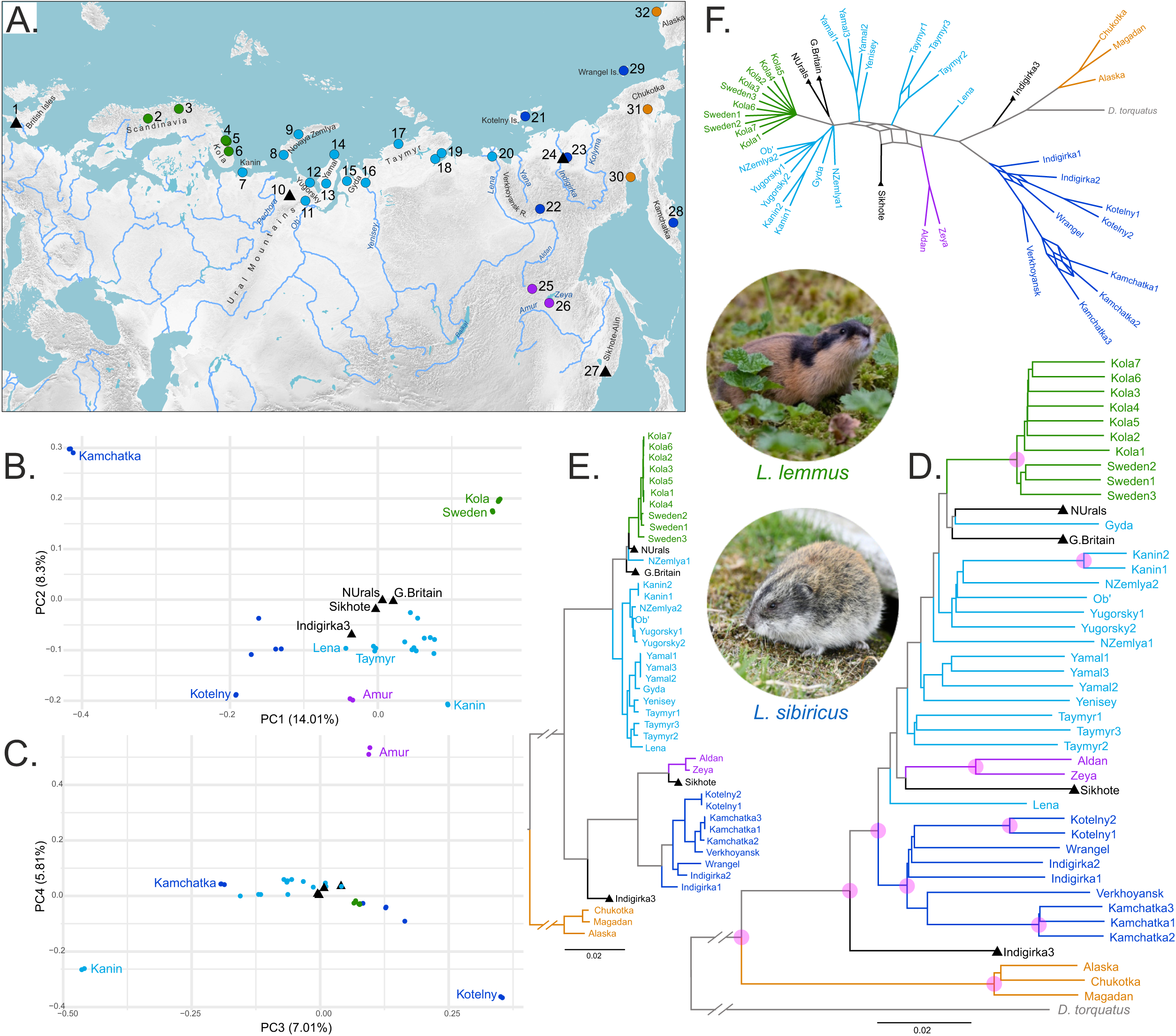
Genetic structure of the genus *Lemmus*. (A) Sampling localities. (B,C) Principal component analysis of the autosomal SNP dataset (*L. trimucronatus* excluded to improve resolution within Palearctic populations). (D) BioNJ tree inferred from autosomal SNPs. Purple circles denote groups with concordance factors >0.9, indicating that these clades were recovered in most phylogenies reconstructed from non-overlapping genomic windows. (E) Maximum-likelihood phylogeny based on complete mitochondrial genomes. All major nodes received maximal bootstrap support. (F) Autosomal phylogenetic network. Colors denote the major groups of the genus *Lemmus* commonly recognized in the literature: *L. trimucronatus* (orange), *L. amurensis* (purple), *L. sibiricus* “east-group” (dark blue), *L. sibiricus* “west-group” (light blue), *L. lemmus* (green), and ancient samples of true lemmings from different time periods (black triangles). Photographs were obtained from https://www.inaturalist.org/.

Two ancient samples presented in this study (Indigirka3, Sikhote) showed strong PMD patterns and moderate endogenous DNA content in raw reads (Fig. S1, Table S2). Two samples (Indigirka1, Indigirka2), although they were found in permafrost ancient layers among mammoth bones, showed no PMD patterns or signs of contamination (Fig. S1). Although there are examples of extracting DNA of very good quality from permafrost (Kistler et al., 2017) and even ancient RNA (Mármol-Sánchez et al., 2026), we chose a conservative approach, therefore treated them as modern samples. Additional sequencing of samples from the same place from the Late Pleistocene can shed light on the authenticity of the samples in question.

The window-based phylogenetic reconstruction and PCA produced largely concordant patterns of genetic structure. In the PCA, most samples clustered within a large group uniting western and eastern populations of *L. sibiricus* together with ancient samples. Several populations were clearly separated from this main cluster, including samples from the Scandinavia, Kola and Kamchatka peninsulas (Fig. 1B), the Kanin Peninsula, the Amur region, and Kotelny Island (Fig. 1C). The same groups also showed high concordance factor values in the window-based phylogenetic analysis (Fig. 1D), strong genetic differentiation. Thus, the groups separated along the first four principal components correspond to the clades with the highest concordance factors in the autosomal tree. The eastern *L. sibiricus* group also showed a high concordance factor, whereas the western group did not. A residual heatmap for the autosomal phylogeny is present in Figure S2. The autosomal phylogenetic network (Fig. 1F) revealed the same major clades as the autosomal tree and additionally demonstrated extensive reticulation within the western *L. sibiricus* group.

The main discrepancy between the PCA and phylogenetic analyses involved the ancient Indigirka3 sample, likely due to differences in the handling of missing genotypes. Whereas the PCA algorithm imputes missing genotypes using mean allele frequencies across the dataset, our pairwise distance approach excludes sites with missing genotypes in either sample, resulting in potentially less biased genetic distance estimates for samples with a high proportion of missing genotypes.

The mitochondrial phylogeny (Fig. 1E) demonstrates the division into western and eastern clusters and the paraphyly of the Siberian lemming. One of the notable discrepancies with the nuclear phylogeny is the position of the sample from Novaya Zemlya Island (NZemlya1), which belongs to the Norway lemming according to mtDNA, and to the Siberian lemming according to autosomal DNA. Another incongruence between the mt and autosomal phylogenies is the position of Amur lemmings (together with the ancient Sikhote sample). While they are sister to the Eastern clade in the mt tree, they are placed within the western *L. sibiricus* clade in the autosomal phylogeny.

### 3.2 Genetic Connectivity Among Palearctic True Lemmings Populations

The optimal number of clusters inferred by LEA was K = 3 (Fig. S4). Two analyses of genetic connectivity among Palearctic lemming populations complemented the results of the genetic structure analyses. Both D-statistics (Fig. 2A) and LEA analyses (Fig. 2B) consistently indicated gene flow between geographically adjacent populations. The most prominent signal involved gene flow between the western and eastern groups of Palearctic lemmings. Samples from the Taymyr Peninsula, Lena River delta, and the Amur region showed particularly strong genetic admixture with the eastern *L. sibiricus* group. Samples from the western *L. sibiricus* group from the lower reaches of the Ob’ River, and the Yugorsky and Gyda peninsulas showed genetic admixture with the Norway lemmings. The results for D-statistics using *Dicrostonyx* as an outgroup are present in Figure S3.

**Figure 2.**
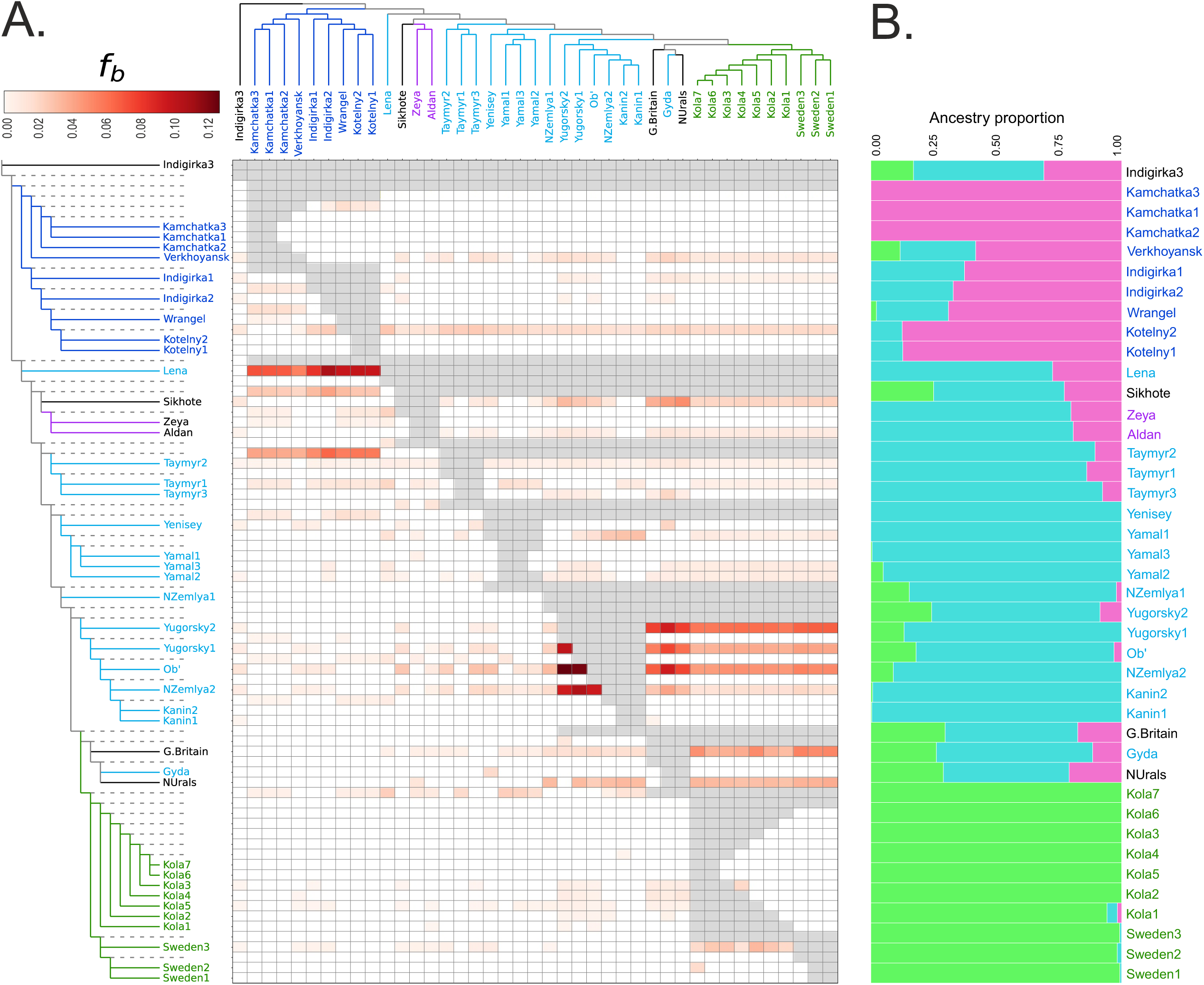
Genetic connectivity within the Palearctic branch of *Lemmus*. (A) The branch-specific statistic *f*b identifies excess sharing of derived alleles between the branches of the phylogeny on the y-axis and the monophyletic groups on the x-axis. The autosomal phylogeny was used as the basis for the branch statistic, and grey cells correspond to tests inconsistent with the underlying topology. (B) Inferred ancestry coefficients for K = 3 ancestral populations.

All ancient samples showed ancestry profiles containing components associated with all major modern genetic lineages in LEA analyses (Fig. 2B). The G.Britain and NUral samples showed a prominent excess of shared alleles with modern Norway lemmings. The Sikhote sample showed genetic affinity to the eastern *L. sibiricus* group in D-statistic analyses, together with modern Amur lemmings. The Indigirka3 sample did not show a clear signal of genetic affinity to any modern group in D-statistic analyses, in contrast to LEA analyses. This discrepancy is likely related to differences in the way the methods handle missing genotypes.

### 3.3 Drivers of Genetic Structure

The Mantel test showed a strong correlation between pairwise geographic and genetic distances. Most pairwise comparisons were distributed close to the regression line (Fig. 3A). As highlighted in Fig. 3B, outliers from the regression (exceeding 2 standard deviations) corresponded to within-group comparisons for populations with low genetic diversity. The same groups also showed the presence of relatively short ROH fragments (100–500 kbp), with the greatest number of ROHs observed in Norway lemmings and in lemmings from the Kanin and Kamchatka peninsulas (Fig. 3C). Genome-wide heterozygosity estimates were also reduced in these groups (Fig. 3D).

**Figure 3.**
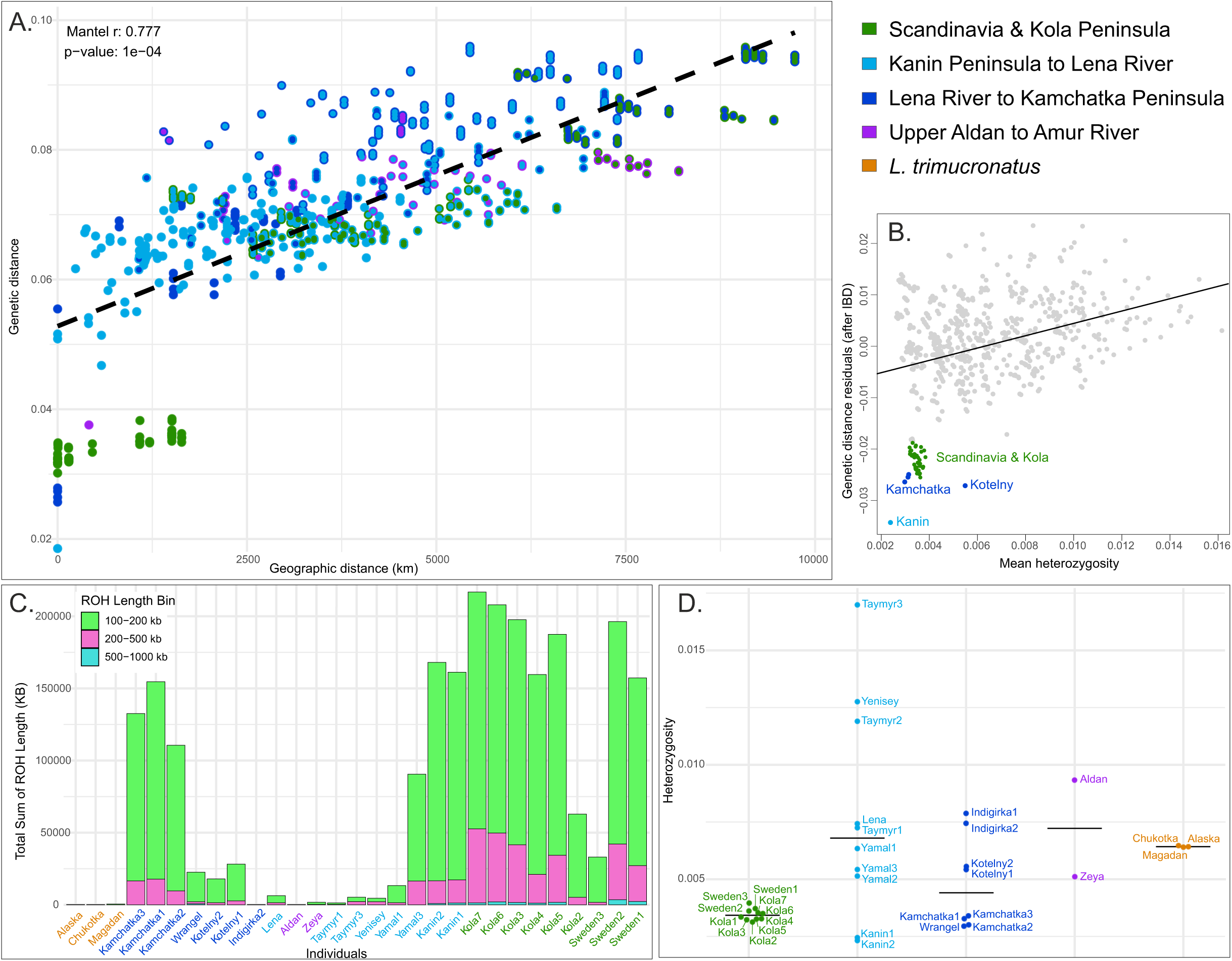
Isolation-by-distance patterns and genomic signatures of bottlenecks in several populations of true lemmings. (A) Regression plot of pairwise genetic distance versus geographic distance. Each point represents a pairwise comparison, with colors indicating the population assignments of the two samples in each pair. Results of the Mantel test are shown in the upper left corner. (B) Residuals from the regression shown in panel A plotted against the mean heterozygosity of each sample pair. Pairwise comparisons highlighted in color deviate from the regression in panel A by more than two standard deviations. (C) Sample-specific ROH distributions. (D) Individual genome-wide heterozygosity estimates.

Interestingly, Amur lemmings did not show signs of inbreeding, as they exhibited moderate heterozygosity and an absence of ROHs. *Lemmus trimucronatus* samples also lacked genomic signatures of inbreeding. The regression of pairwise geographic and genetic distances including *L. trimucronatus* samples (Fig. S4) clearly separated two groups, as all comparisons between the Palearctic and Nearctic branches showed substantially greater genetic distances relative to geographic distance.

### 3.4 Demographic History of Norway Lemmings

The final dataset used for demographic inference contained 246 million called sites shared across all samples after depth and missing-data filtering, including 2.5 million variable sites. The best CLAIC-supported model inferred with GADMA (Fig. 4) indicates a bottleneck in the history of Norway lemmings that occurred shortly before the split between the Scandinavian and Kola Peninsula populations. The onset of the bottleneck was inferred at 12 ka BP. The best model according to log-likelihood recovered a similar bottleneck in the history of Norway lemmings preceding the population split at the time corresponding to the Pleistocene–Holocene transition (Fig. S6). This concordance across alternative model structures suggests that the bottleneck signal represents a robust feature of the demographic history inferred from the data.

**Figure 4.**
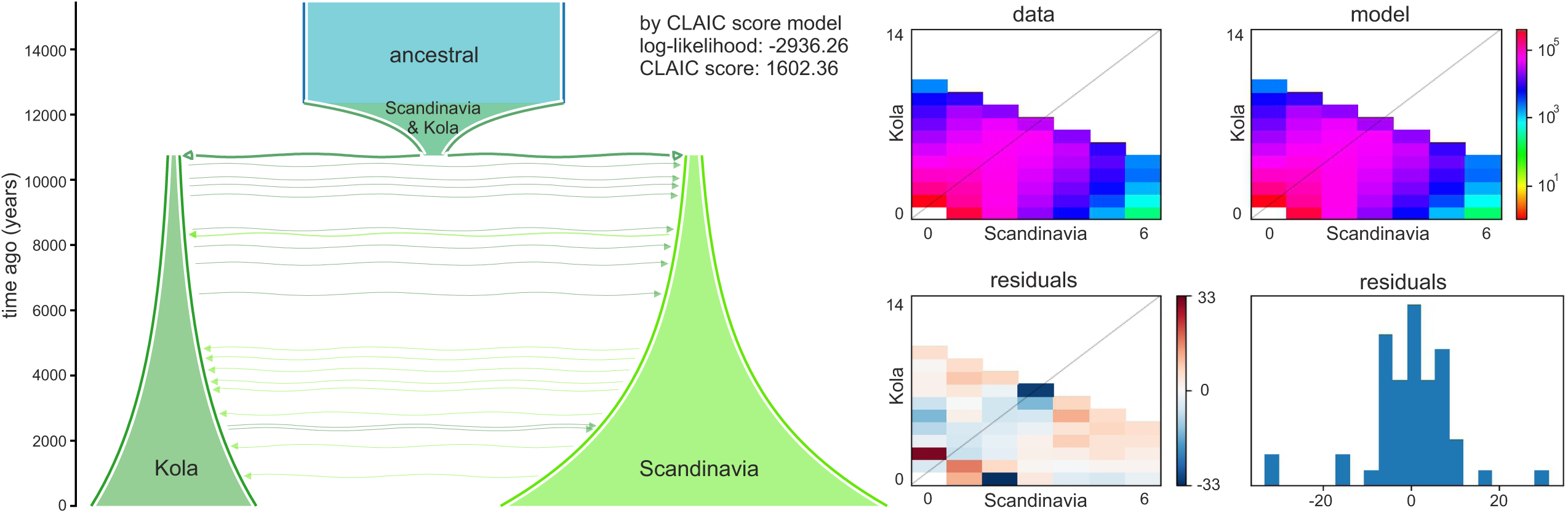
Demographic history of Norway lemmings inferred using GADMA. The left panel shows the best-supported demographic model by CLAIC. The upper right panel shows the observed folded site frequency spectrum (SFS) and the modelled SFS. The lower right panel shows residuals between the observed and modelled SFS.

Confidence intervals estimated from bootstrapped SFS replicates for the best CLAIC-supported model did not always encompass the parameter estimates of the model itself and in some cases were closer to the estimates recovered by the best log-likelihood model (Table S3). This discrepancy may be related to the small sample size and uneven representation of the populations in the dataset. Nevertheless, the confidence intervals consistently supported a bottleneck preceding population splitting at approximately the time of the Pleistocene–Holocene transition. Therefore, we consider this bottleneck signal to be the most robust result of the demographic analyses, whereas the remaining demographic parameters should be interpreted with caution and require further investigation.

## 4 Discussion

### 4.1 Genetic Structure of the Palearctic *Lemmus* Mirrors the Post-LGM Range Shifts

The modern genetic structure of Palearctic true lemmings reflects the superposition of various evolutionary processes acting at different temporal scales. Analyses of nuclear genome diversity provided here together with the information on mt genome analyses (Lord et al., 2025; Panitsina et al., 2026) and paleogeographic data (Markova et al., 2019, 1995; Tiunov and Panasenko, 2011) allow us to partially reconstruct the evolutionary history of the Palearctic branch of the genus in the Late Pleistocene.

Based on the WGS data we showed that genome structure of the Palearctic branch of the *Lemmus* genus consists of the interconnected populations inhabiting tundra from Novaya Zemlya Island on the west, to the Kolyma River on the east, near that presumable border between Nearctic and Palearctic branch is located. Given the central geographic position of this population within the Palearctic *Lemmus* range, we will refer to this group as central. Inside the central group we detect weak signal of genetic separation onto western and eastern clades, corresponding to the deep mt divergence observed previously.

At the same time, several geographically isolated peripheral populations are clearly genetically separated from this central group. These include the Norway lemming (Scandinavia and Kola Peninsula) population, the lemmings from the eastern shore of Kamchatka Peninsula, the Kotelny Island population, the Kanin Peninsula population, and the Amur lemmings (South-East Siberia). Most of these populations are currently isolated from the central group. Next through the text we will refer to them as peripheral.

Although several peripheral populations form monophyletic clusters with high concordance factors in the autosome phylogeny, they remain embedded within the broader continuum of genetic variation of the central group. The autosomal phylogeny shows a largely comb-like branching structure, with geographically peripheral populations occupying terminal positions within this continuum rather than forming deeply separated lineages. The overall nuclear divergence of peripheral populations from neighbouring ones largely follows the general isolation-by-distance relationship (Fig. 3A). Together, these patterns suggest that the strong phylogenetic concordance observed in peripheral groups does not primarily reflect long-term independent evolution leading to the acquiring of unique adaptation features through mutation processes and subsequent ecological divergence as well as reproductive isolation. Instead, we interpret it as a consequence of demographic processes accompanying geographic isolation at the range margins, particularly bottlenecks and enhanced genetic drift, which promoted local lineage sorting in the autosomal genome.

This interpretation is further supported by reduced heterozygosity and the presence of ROHs in all peripheral populations except the Amur lemmings. The predominance of relatively short ROH fragments suggests bottlenecks of moderate intensity that occurred in the more distant past (Ceballos et al., 2018). For Norway lemmings, demographic modeling with GADMA (Fig. 4) also supported a bottleneck scenario, with the strongest reduction in effective population size occurring after the LGM. We hypothesize that the isolation of these peripheral populations was established during the Late Glacial and early Holocene, following the LGM. Founder effects during postglacial colonization, followed by prolonged isolation and reduced effective population sizes, could have generated the observed genetic patterns. This scenario is consistent with mt divergence estimates (Lord et al., 2025; Panitsina et al., 2026) for most peripheral lineages, as well as with paleogeographic evidence indicating that the broad LGM distribution of lemmings subsequently contracted and shifted northward. Moreover, ancient LGM (Sikhote) and post-LGM (G.Britain, N.Urals) samples analysed here showed strong affinity to the central group of modern lemmings, according to all analyses. Given their broad range of sampling localities from the British Isles to Sikhote-Alin Mountains this supports the view of much more connected lemmings population during LGM and Pleistocene-Holocene transition period.

The mechanisms underlying isolation likely differed among populations. Isolation of populations of Scandinavia and Kola peninsulas and the Kotelny Island was probably associated with postglacial sea-level rise, whereas the Kamchatka and Amur populations were separated from the central range by extensive areas of taiga biome, where true lemmings do not occur due lose in competition with other vole species abundant in taiga (Ehrich et al., 2020; Rausch and Rausch, 1975). The causes of isolation in the Kanin Peninsula population remain unclear, since this region is not presently separated from the Siberian mainland by obvious unsuitable habitat. However, only two individuals from the Kanin Peninsula were analyzed in this study, and precise locality data for these samples are unavailable.The isolated population of lemmings on Wrangel Island requires additional sampling to assess the level of genomic divergence.

The absence of clear signs of bottlenecks in the Amur lemmings is consistent with the view of this population as a remnant of the broad LGM lemming range (Markova et al., 2019). Signals of gene flow with other lemming populations further support former connectivity between Amur lemmings and other ancient populations. Moreover, our ancient sample from Sikhote-Alin (dated to 18.5 ka BP) does not reveal genomic separation from the central group, indicating that lemmings from the Russian Far East remained connected with other populations during the LGM. Although the deep divergence of the Amur mt lineage could suggest a more ancient separation, the lack of pronounced nuclear divergence indicates that mtDNA may retain signatures of older demographic processes that are no longer detectable in the autosomal genome. Because mtDNA is maternally inherited, it has a lower effective population size and therefore undergoes lineage sorting faster than the nuclear genome, leading to a stronger effect of genetic drift. In addition, mt lineages do not recombine, allowing deeply divergent haplotypes to persist for long periods of time.

For Norway lemmings, the hypothesis of pre-LGM isolation and persistence within a Scandinavian refugium during the LGM was previously proposed based on mtDNA analysis (Lagerholm et al., 2014) and whole-genome data (Feinauer et al., 2026). The revealed pattern in the nuclear genome here can be consistent with both pre-LGM isolation and subsequent persistence in refugia and post-LGM colonisation, as both of these scenarios lead to the bottleneck. Although mt divergence times estimation for modern Norway lemmings repeatedly testifies against Norwegian refugium, as estimates based on tip dating have yielded LGM or post-LGM divergence time. We consider tip dating calibration as the most appropriate one for evolutionary recent events (Duchêne et al., 2014; Ho et al., 2005).

In Lord et al. (2025), another hypothesis of pre-LGM separation of Norway lemmings was proposed. The authors expanded the Norway lemming group by including two post-LGM ancient samples from the British Isles and Northern Urals. These ancient samples occupied a basal position relative to the modern Norway lemming clade in both mt and nuclear phylogenies. Inclusion of these samples increased the inferred mt divergence time of the expanded Norway lemming clade to approximately 38 ka BP. As discussed in Panitsina et al. (2026), this hypothesis would imply long-term genetic isolation of lemmings distributed from the British Isles to the Ural region from other Eurasian populations throughout much of the Late Pleistocene. That appears inconsistent with the paleogeographic evidence discussed above. Moreover, our autosomal analyses demonstrate clear genetic affinity of these ancient samples to the central group, providing additional evidence against the proposed scenario.

A more complex pattern is observed within the central group itself, particularly in the relationship between the western and eastern clades of *L. sibiricus*. Lemmings inhabiting the territory east of Lena River form a highly concordant monophyletic clade across genomic windows, suggesting a period of historical isolation. However, admixture and D-statistics analyses simultaneously demonstrate substantial gene flow between the eastern and western clades. Mitochondrial analyses indicate that these lineages diverged during the Late Pleistocene, before the LGM. We therefore hypothesize that western and eastern Siberian lemmings persisted as partially isolated populations during part of the Late Pleistocene, accumulating nuclear and mt divergence, although the reasons for this isolation remain unclear. During the LGM, expansion of suitable tundra habitats may have reconnected these populations, leading to secondary contact, gene flow, and partial erosion of previously accumulated genetic differences. Under this scenario, the eastern lineage retained part of its divergence, which today appears as strong phylogenetic concordance, whereas admixture reduced overall nuclear differentiation between the eastern and western groups. This interpretation is consistent with several observations. First, differentiation between eastern and western clades of lemmings is weaker than that observed in isolated peripheral populations, as these groups are not clearly separated along the major principal components (Fig. 1B,C). Second, pairwise comparisons between eastern and western populations do not form strong outliers in the isolation-by-distance analysis. Together, these patterns suggest incomplete divergence accompanied by substantial historical gene flow rather than long-term independent evolution.

In general, our results suggest that the modern genetic structure of Palearctic lemmings was shaped largely during the last major climatic cycle, including the LGM and the Pleistocene–Holocene transition. Given that the evolutionary history of the genus spans at least 2 million years (Louis et al., 2024), the present-day genetic structure represents only a small fragment of its vast evolutionary history. Our Indigirka3 sample, dated to approximately 40 ka BP (Lopatin et al., 2019), provides a glimpse into pre-LGM diversity. This specimen is genetically distinct from all other analyzed Palearctic lemmings and may represent a Late Pleistocene population from northeastern Siberia that did not persist into the present day. The reasons for this population turnover remain beyond the scope of the present study.

### 4.2 Species Delimitation Within True Lemmings

The species status of a given taxon is a complex issue, with more than 26 proposed species concepts and definitions (De Queiroz, 2007). Moreover, species delimitation may also be influenced by conservation policy and legal considerations (Frankham et al., 2012). Therefore, any discussion of species status necessarily depends on the particular species concept applied. As this work provides only phylogenomic results, we discuss here the problem of taxonomic separation within true lemmings from the perspective of the Phylogenetic Species Concept.

All analyses consistently revealed deep genetic separation between the Palearctic and Nearctic branches. The two groups were clearly separated in the nuclear phylogeny, and pairwise comparisons between them showed substantially greater genetic distances relative to geographic distance than comparisons within either branch (Fig. S5). These findings are consistent with previously reported karyotypic differences and partial reproductive isolation (Gileva et al., 1984; Pokrovski et al., 1984). Although D-statistics detected signals of gene flow between the branches (Fig. S3), the timing and evolutionary significance of this gene flow remain unclear.

In contrast, we did not find any clear separation into distinct genetic groups within the Palearctic lemmings. The comb-like pattern observed in the Palearctic clade on the phylogenetic tree provides insights into the evolutionary processes affecting their genomes, but it also has taxonomic implications. When assigning or maintaining species status for any peripheral group we described here, one must consider that all peripheral populations are nested within the central group of lemmings in the species tree (i.e., the autosomal phylogeny). This means that defining any of these groups as separate species would automatically result in the creation of a paraphyletic taxon. For example, retaining species status for the Norway lemming would render *Lemmus sibiricus* paraphyletic, as Norway lemmings are among the descendants of the most recent common ancestor of western clade of the central group.

The alternative possibility of splitting Palearctic lemmings into only two species corresponding to the western (considering *L. amurensis* and western lineage of *L. sibiricus* as junior synonyms of *L. lemmus*) and eastern (*L. sibiricus* eastern clade) clades of the central group, including the peripheral populations within them, is also unsupported from our point of view. The genetic differentiation between eastern and western clades is relatively weak even on the scale of the Palearctic branch. The strongest separation on PCA is observed between peripheral populations along the first principal components. In addition, we demonstrated substantial gene flow between groups, while the observed genetic distances appear to be shaped primarily by geographic distance as inferred by IBD analyses.

### 4.3 Data Quality and Robustness of Genomic Patterns

We believe that the genomic patterns described above are robust to potential biases caused by uneven sequencing coverage across samples. Owing to the nature of the material, which included ancient and museum specimens, such variation in coverage was unavoidable. However, during bioinformatic processing we carefully controlled for the possible clustering of low-coverage samples caused by high levels of missing data. In some analyses we nevertheless observed contradictory patterns associated with the high missingness of certain ancient samples, such as the inconsistent position of the ancient Indigirka3 specimen in the PCA and autosomal phylogeny. In these cases, we interpreted the results cautiously and relied primarily on patterns that were concordant across multiple analytical approaches. Overall, we believe that this study represents the first comprehensive investigation of genome-wide structure in the genus *Lemmus*, with sampling spanning nearly its entire Palearctic range.

### 4.4 Regional Variation in Post-Glacial Diversity

The fate of lemmings during the drastic climatic changes of the Pleistocene–Holocene transition, investigated here, allows us to discuss the broader concept of Arctic species population dynamics. In Stojak and Jędrzejewska (2022), it was proposed that Arctic species lost much of their genetic diversity during climatic cycles because of the absence of continuous refugia in which distinct genetic lineages could persist.

Here, however, we show that although the European population of lemmings, which descendants are represented today by Norway lemmings, indeed lost much of its genetic diversity during the Pleistocene–Holocene transition, lemmings from the Asian part of the genus range still retain substantial genomic diversity. We suggest that a similar pattern may occur in other Arctic species and that understanding their evolutionary response to climate change is therefore impossible without extensive sampling across Russian Siberia.

## Author Contributions

N.A. and I.D. conceived the study. I.D. performed the research with input from S.B. and V.P. and substantial visualisation support from T.P. Authors: N.S., A.P., A.K., M.T., A.L., L.L. and N.A. contributed samples and/or participated in sampling campaigns. I.D. wrote the manuscript, with contributions from N.A., T.P. and L.L. Funding acquisition: N.A. All authors provided feedback on the manuscript and approved the final revised version.

## Acknowledgements

The authors thank all colleagues who helped in numerous field research trips and shared materials necessary for the study. We thank the curators of the scientific fund collection of mammals Makarova O.V. and Maksimova E.R. for invaluable help in sample selection. The financial support for the study for I.D., T.P., V.P., S.B. and N.A. was provided by the Russian Science Foundation grant N19-74-20110-P. For M.T. the research was carried out within the state assignment of the Ministry of Science and Higher Education of the Russian Federation (theme No. 124012200182-1).

## Funding

This research was funded by the Russian Science Foundation grant N19-74-20110-P.

## Ethics Statement

This study did not require official or institutional ethical approval.

## Conflicts of Interest

The authors declare no conflicts of interest.

## Data Availability Statement

Raw sequence reads have been deposited in the NCBI Sequence Read Archive under BioProject accession PRJNA590630.

## Code availability

Scripts used to raw sequencing data processing and subsequent analyses in this study are available at https://github.com/VanoBorodatovich/WGS-analyses. Script to create block-bootstrap of VCF file for subsequent GADMA analyses is available at https://github.com/VanoBorodatovich/Block_bootstrap_for_vcf.

## Benefit-sharing statement

This research was conceptualized, funded, and executed entirely by scientists from Russian research institutions, utilizing modern and fossil lemming specimens originating within the territory of the Russian Federation. Consequently, benefits from this work are shared directly through national scientific capacity building and the public availability of all primary data, which contribute to the global understanding of Quaternary Arctic biodiversity.

## Notes

### Competing Interest Statement

The authors have declared no competing interest.

