## Supplementary for "A Whole-Genome and Ancient DNA Perspective on the Drivers of Genetic Diversity and Structure in Palearctic True Lemmings"

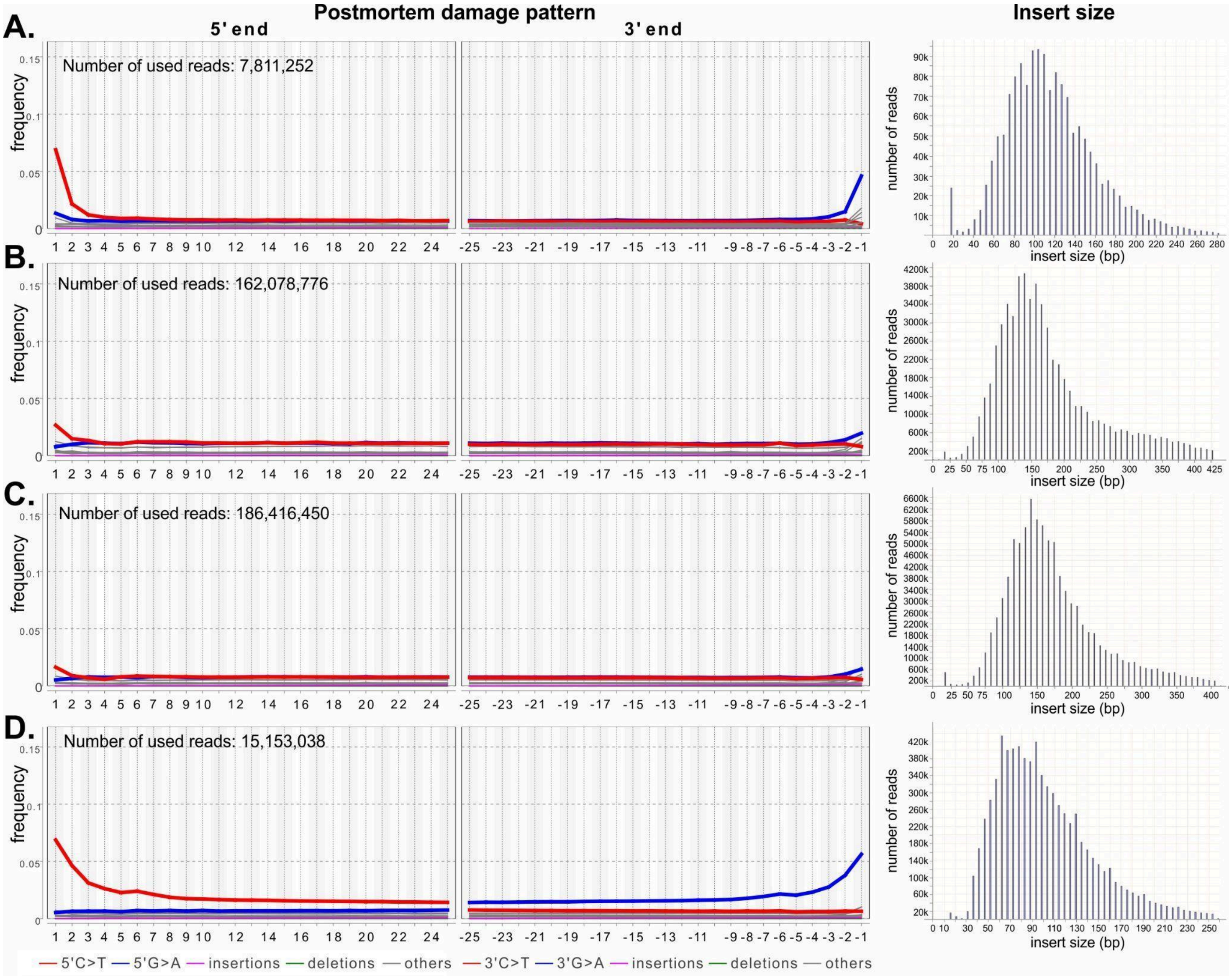

**Figure S1**. Ancient DNA damage patterns. (**A**) *Lemmus amurensis*, Perspektivnaya, 18.550 kya; (**B**) *Lemmus sibiricus*, Ogorokha, 28.390 kya; (**C**) *Lemmus sibiricus*, Ogorokha, 18.972 kya; (**D**) *Lemmus sp*., Tirekhtyakh, 41.595 kya. The nucleotide positions of the 5' and 3' ends of the reads are shown on the horizontal axis, with the proportion of incorrectly inserted bases represented on the vertical axis. The red and blue colors indicate cytosine-to-thymine substitution and guanine-to-adenine substitution, respectively.

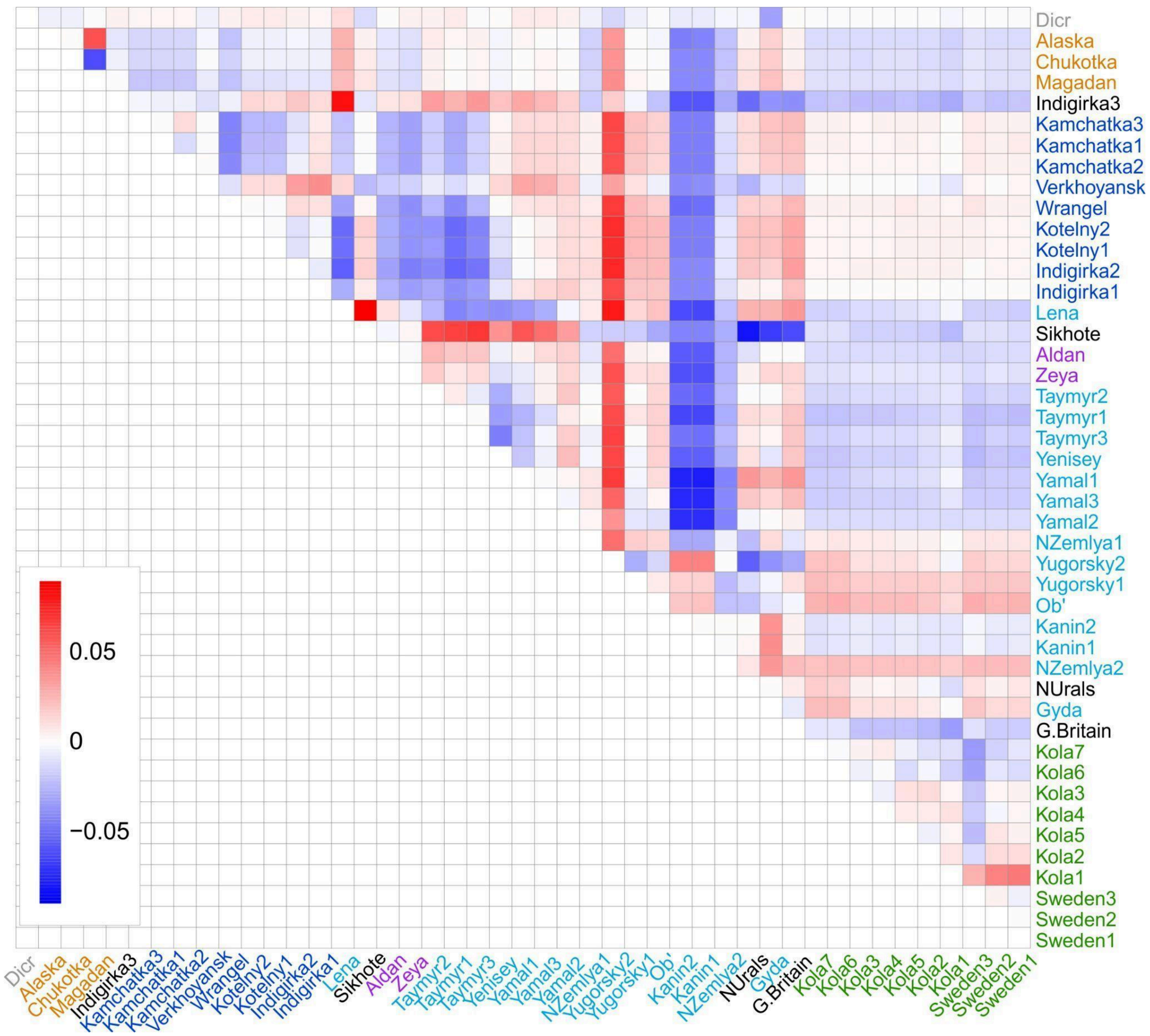

**Figure S2**. Residual heatmap for the autosomal dendrogram (Fig. 1D), showing differences between path lengths in the dendrogram and actual genetic distances in the underlying distance matrix. Blue ("attraction") indicates sample pairs that are more similar than suggested by the dendrogram, whereas red ("repulsion") indicates the opposite.

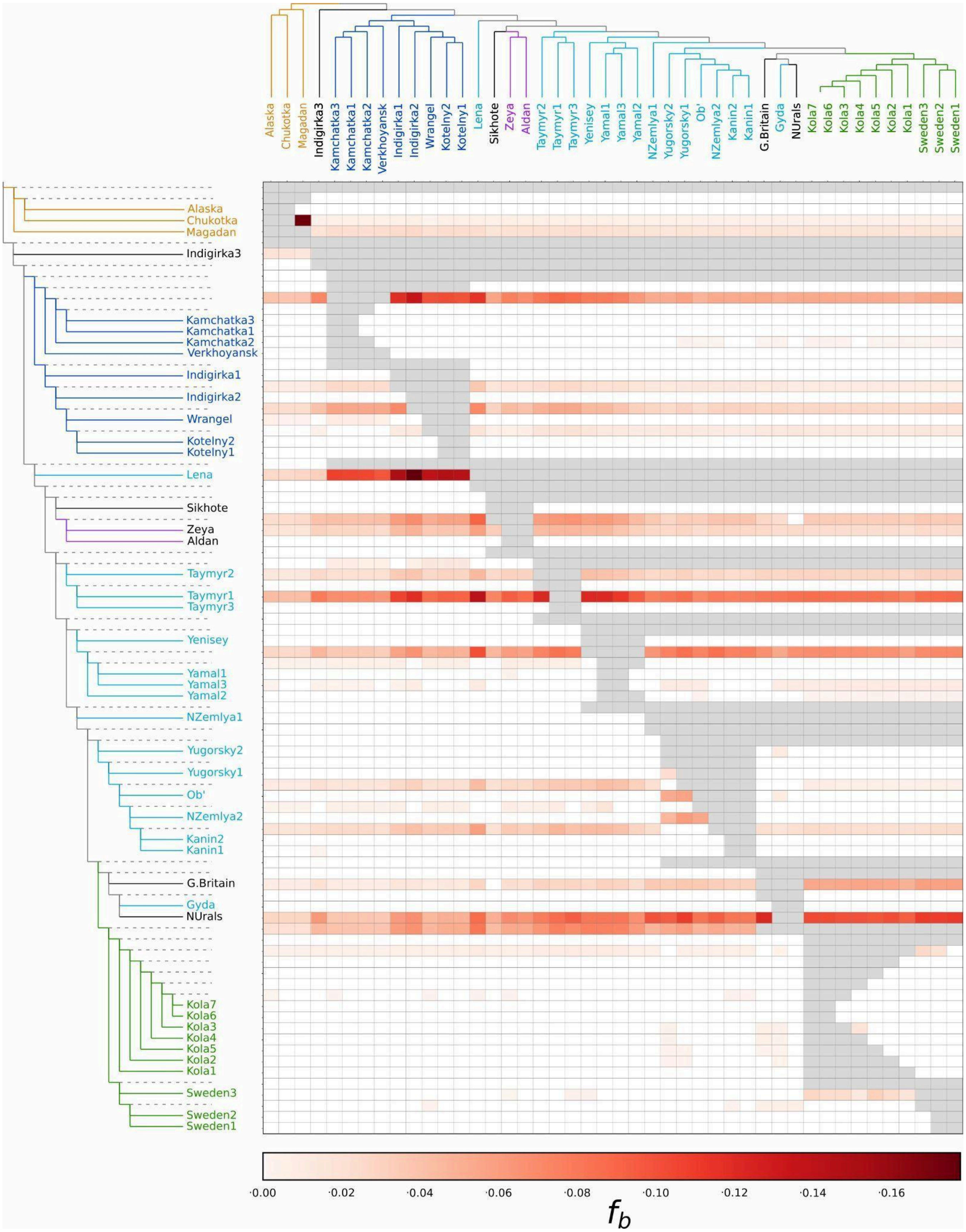

**Figure S3.** The branch-specific statistic *f*b identifies excess sharing of derived alleles between the branches of the phylogeny on the y-axis and the monophyletic groups on the x-axis. The autosomal phylogeny including *Dicrostonyx* as an outgroup was used as the basis for the branch statistic, and grey cells correspond to tests inconsistent with the underlying topology.

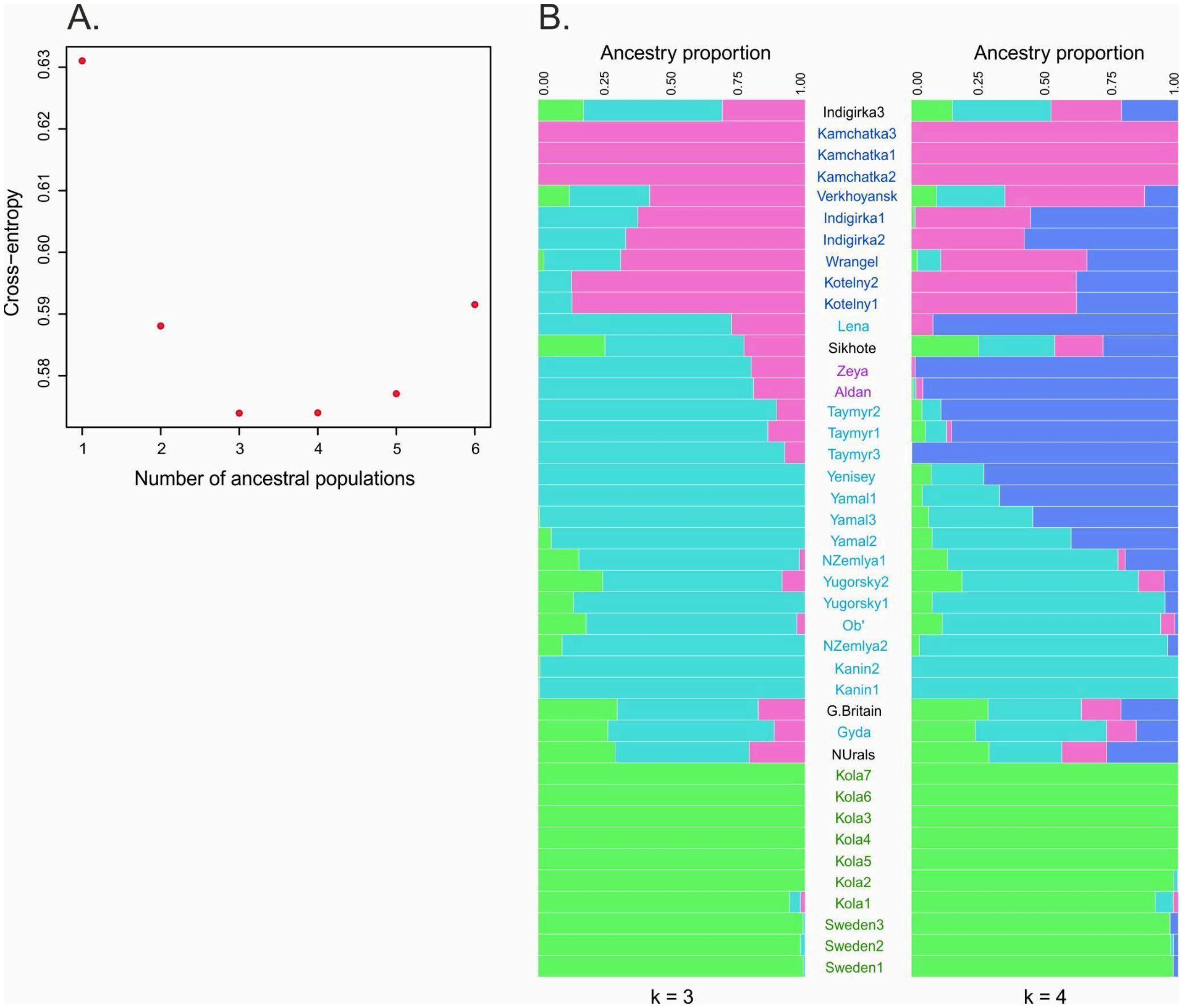

**Figure S4**. (A) The cross-entropy plot for the number of clusters (K = 2–6). (B) Inferred ancestry coefficients for K = 3 and K = 4 ancestral populations.

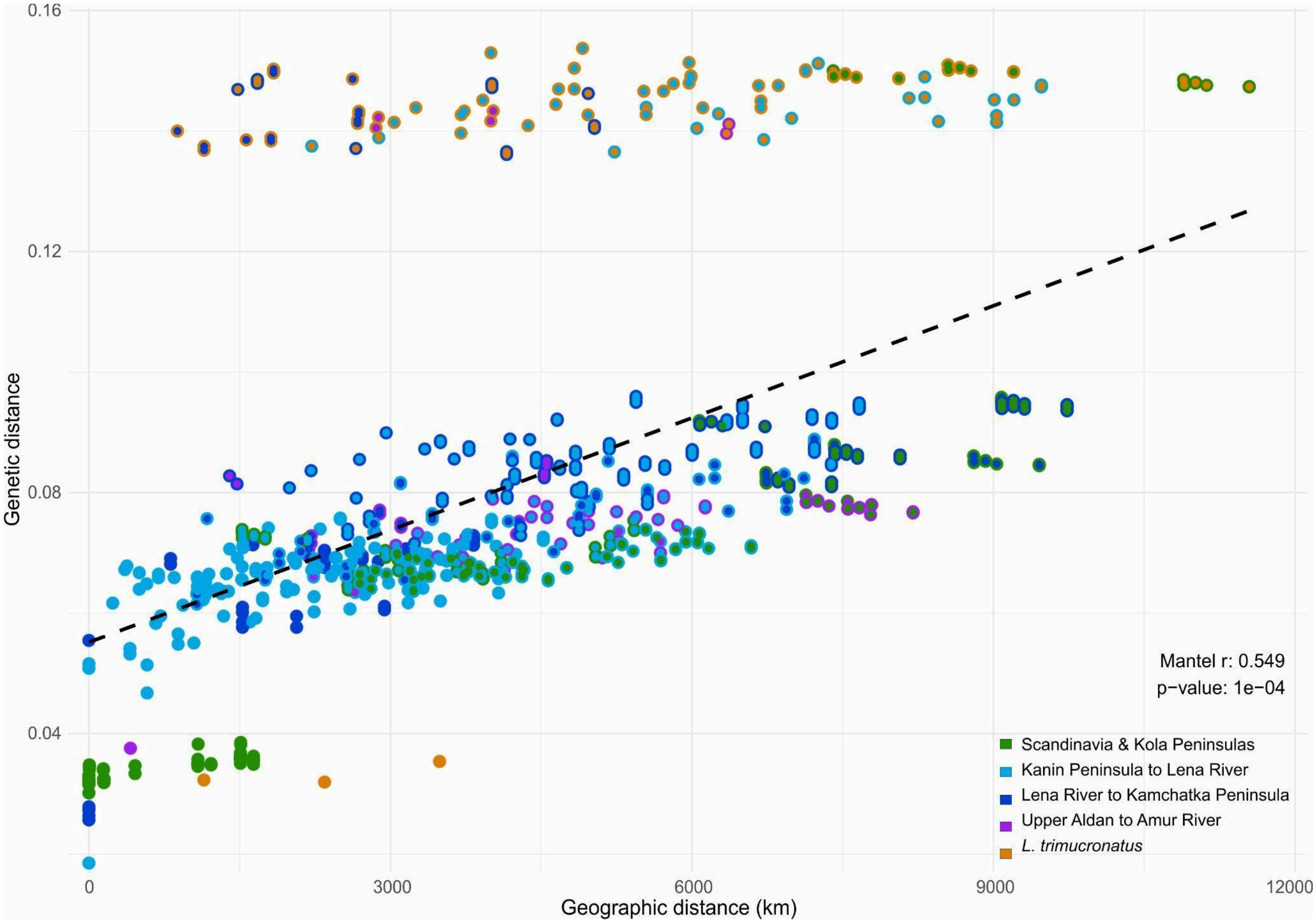

**Figure S5**. Regression plot of pairwise genetic distance versus geographic distance. Each point represents a pairwise comparison, with colors indicating the population assignments of the two samples in each pair. Results of the Mantel test are shown in the lower right corner.

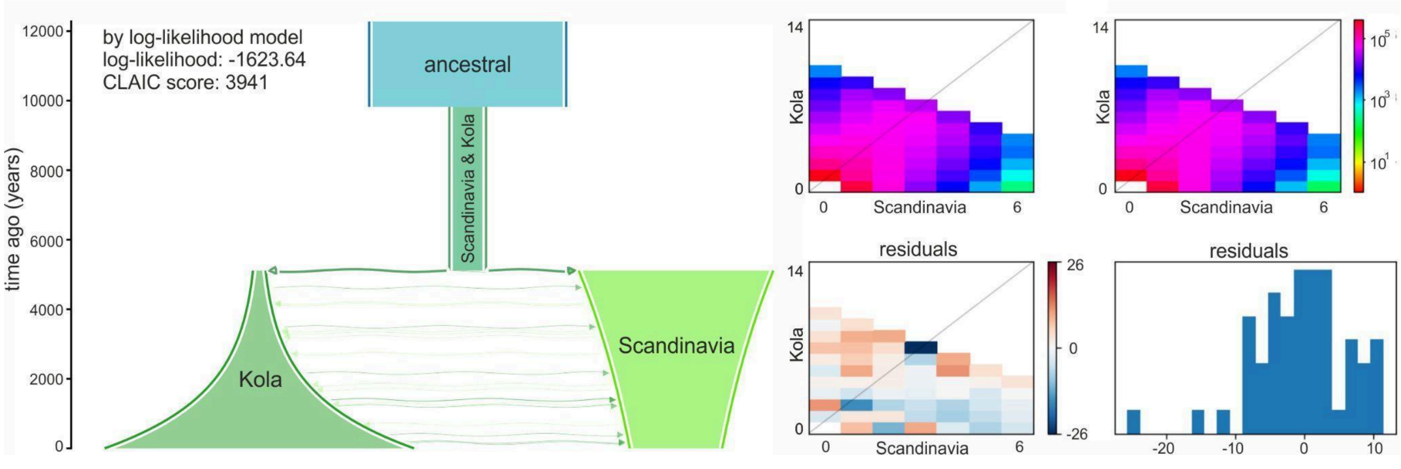

**Figure S6**. Demographic history of Norway lemmings inferred using GADMA. The left panel shows the best-supported demographic model by log-likelihood. The upper right panel shows the observed folded site frequency spectrum (SFS) and the modelled SFS. The lower right panel shows residuals between the observed and modelled SFS.

### Table S1. Materials used in the study

| Map ID | Species | Locality | Lat | Lon | Sample name | Voucher ID | Tissue ID | Tissue type | Sex | Reference |
|---|---|---|---|---|---|---|---|---|---|---|
| 1 | *L. lemmus* | England, Bridged Pot | 51.23 | -2.68 | G.Britain | | BP13k | ancient | m | Lord et al., 2025 |
| 2 | | Sweden, N Jämtland, Gussvattnet | 64.46 | 14.3 | Sweden1 | | LEM01 | modern | m | |
| | | | | | Sweden2 | | LEM02 | modern | f | |
| 3 | | Sweden, Northern Lappland, Riksgränsen | 68 | 18 | Sweden3 | | LEM05 | modern | m | |
| 4 | | Russia, Murmansk Region, Kola Peninsula, Tumanny | 68.98 | 35.75 | Kola1 | ZIN 86357 | 393 | modern | f | current study |
| 5 | | Russia, Murmansk Region, Kola Peninsula, Teriberka | 69.013 | 35.707 | Kola2 | | 6290 | modern | f | current study |
| | | | | | Kola3 | | 6291 | modern | m | current study |
| | | | | | Kola4 | | 6293 | modern | m | current study |
| | | | | | Kola5 | | 6294 | modern | m | current study |
| 6 | | Russia, Murmansk Region, Kola Peninsula, Varzina river | 68.18 | 38.06 | Kola6 | | LEM03 | modern | m | Lord et al., 2025 |
| | | | | | Kola7 | | LEM04 | modern | m | |
| 7 | *L. sibiricus* | Russia, Nenets AO, Kanin Peninsula | 67 | 45 | Kanin1 | | SIW08 | modern | f | |
| | | | | | Kanin2 | | SIW09 | modern | m | |
| 8 | | Russia, Novaya Zemlya, Propashchaya Bay | 71.1458 | 53.793 | NZemlya1 | ZIN 11037 | 6350 | museum | f | current study |
| 9 | | Russia, Novaya Zemlya, Krestovaya Bay | 74.11 | 55.4 | NZemlya2 | ZIN 22442 | 6326 | museum | m | current study |
| 10 | | Russia, Nenets AO, Pymva Shor | 67.1 | 60.51 | NUrals | | PS10k | ancient | m | Lord et al., 2025 |
| 11 | | Russia, Yamalo-Nenets AO, upper Sob' River (tributary of Ob' River) | 67.0615 | 65.4839 | Ob' | ZIN 17703 | 6357 | museum | f | current study |
| 12 | | Russia, Nenets AO, Yugorsky Peninsula, the eastern shore of the Kaskaya Bay | 69.2564 | 65.0316 | Yugorsky1 | ZIN 24399 | 6359 | museum | m | current study |
| | | | | | Yugorsky2 | ZIN 24395 | 6360 | museum | f | current study |
| 13 | | Russia, Yamal, Yasaveyto Lake | 69.7439 | 70.0908 | Yamal1 | | 662 | modern | f | current study |
| 14 | | Russia, Yamal, Bely Island, polar station | 73.3311 | 70.0639 | Yamal2 | | 391 | modern | f | current study |
| | | | | | Yamal3 | | 661 | modern | f | current study |
| 15 | | Russia, Yamalo-Nenets AO, Gyda Peninsula, Yuribey River | 70.586 | 76.4064 | Gyda | ZIN 17495 | 6328 | museum | f | current study |

| | | | | | | | | | | |
|---|---|---|---|---|---|---|---|---|---|---|
| 16 | | Russia, Krasnoyarsky Kray, Deryabinsky Island, Yenisey Estuary | 70.7906 | 82.5538 | Yenisey | ZIN 15930 | 6330 | museum | m | current study |
| 17 | | Russia, Krasnoyarsky Kray, North-Western Taymyr, Tolevaya River | 75.6697 | 93.1506 | Taymyr1 | ZIN 81385 | 6336 | museum | f | current study |
| 18 | | Russia, Krasnoyarsky Kray, Taymyr, Near mouth of Bolshaya Balakhna River | 73.6496 | 106.6005 | Taymyr2 | ZIN 25141 | 6337 | museum | m | current study |
| 19 | | Russia, Krasnoyarsky Kray, Eastern Taymyr, Labaznaya River | 74.2395 | 109.2334 | Taymyr3 | ZIN 25151 | 6332 | museum | m | current study |
| 20 | | Russia, Yakutia, Bulunsky District, Lena Delta, Samoylovsky Island | 72.3791 | 126.4842 | Lena | ZIN 86348 | 369 | modern | m | current study |
| 21 | | Russia, Yakutia, Bulunsky District, Kotelny Island, Taas-Ary Peninsula | 75.0355 | 143.4443 | Kotelny1 | | 4991 | modern | f | current study |
| | | | | | Kotelny2 | | 4992 | modern | f | current study |
| 22 | | Russia, Yakutia, Verkhoyansky District, Nelgese River | 64.4469 | 134.0517 | Verkhoyansk | ZIN 16754 | 4464 | museum | m | current study |
| 23 | | Russia, Yakutia, Abyysky District, Middle Indigirka R., Ogoroha R., 54 km SE from Belaya Gora | 68.1854 | 147.0961 | Indigirka1 | | 6346 | ancient | m | current study |
| | | | | | Indigirka2 | | 6347 | ancient | f | current study |
| 28 | | Russia, Kamchatka Peninsula, Uzon | 54.4733 | 160.018 | Kamchatka1 | ZIN 85586 | 396 | modern | m | current study |
| 28 | | | | | Kamchatka2 | ZIN 85587 | 397 | modern | f | current study |
| 28 | | | | | Kamchatka3 | ZIN 85585 | 398 | modern | m | current study |
| 29 | | Russia, Wrangel Island | 71.23 | -179.4 | Wrangel | | SIE10 | modern | f | Lord et al., 2025 |
| 24 | *Lemmus sp.* | Russia, Yakutia, Abyysky District, Middle Indigirka R., Tirekhtyakh R. | 68.5413 | 147.1238 | Indigirka3 | PIN 5663/1 | 6380 | ancient | m | current study |
| 25 | *L. amurensis* | Russia, Yakutia, Neryungri District, Nagorny | 55.95 | 124.91 | Aldan | ZMMU s-150535 | 4576 | museum | f | current study |
| 26 | | Russia, Amur Region, Zeysky District, Pikan | 53.6771 | 127.4628 | Zeya | ZMMU s-91378 | 4568 | museum | f | current study |
| 27 | | Russia, Primorsky Kray, Levaya Ilistaya R., Perspektivnaya cave | 43.6638 | 132.7274 | Sikhote | | 6320 | ancient | f | current study |
| 30 | *L. trimucronatus* | Russia, Magadan Region, Gizhiga Bay | 61.9153 | 159.2765 | Magadan | | 4541 | modern | f | current study |
| 31 | | Russia, Chukotka, 150 km N from Anadyr, Tanurer R. | 66.1547 | 175.7622 | Chukotka | ZIN 93413 | 377 | modern | m | current study |
| 32 | | USA, Alaska, Barrow | 71.29 | -156.78 | Alaska | | TRI11 | modern | m | Lord et al., 2025 |
| n/a | *Dicrostonyx torquatus* | Russia, Yakutia, Bulunsky District, Lena Delta, Samoylovsky Island | 72.3791 | 126.4842 | Dicr | | 371 | modern | f | current study |

**Table S2. Summary statistics of raw WGS data and alignment. Mean coverage was calculated after all filtering steps, with overlapping bases from paired-end reads counted only once.**

| Map ID | Sample name | Tissue ID | Tissue type | M pairs of raw reads sequenced | GC | % of target DNA if contaminated | % bp trimmed | M Pairs aligned | Gen error rate | Mean insertion length and sd | % of duplicates | Mean coverge | Autosome heterozygosity |
|---|---|---|---|---|---|---|---|---|---|---|---|---|---|
| 1 | G.Britain | BP13k | ancient | 380.4 | 58% | 2% | 60.88% | 6.9 | 0.79% | 50/15 | 25.50% | 0.1 | NA |
| 2 | Sweden1 | LEM01 | modern | 226.4 | 41% | None | 9.76% | 221.7 | 0.82% | 358/88 | 12.70% | 19.05 | 0.00376096 |
| | Sweden2 | LEM02 | modern | 216.8 | 41% | None | 9.82% | 212.0 | 0.83% | 377/91 | 11.10% | 18.99 | 0.00364968 |
| 3 | Sweden3 | LEM05 | modern | 268.2 | 40% | None | 4.55% | 261.9 | 0.67% | 161/29 | 11.20% | 15.67 | 0.00401257 |
| 4 | Kola1 | 393 | modern | 42.7 | 41% | None | 3.92% | 41.4 | 0.73% | 258/49 | 12.10% | 3.73 | 0.00349966 |
| 5 | Kola2 | 6290 | modern | 63.1 | 41% | None | 5.21% | 61.6 | 0.76% | 299/72 | 6.20% | 5.97 | 0.00297824 |
| | Kola3 | 6291 | modern | 80.1 | 41% | None | 4.80% | 78.5 | 0.76% | 288/70 | 6.20% | 7.45 | 0.00316806 |
| | Kola4 | 6293 | modern | 77.2 | 41% | None | 7.28% | 75.2 | 0.83% | 344/89 | 8.00% | 7.42 | 0.00320524 |
| | Kola5 | 6294 | modern | 80.1 | 41% | None | 5.40% | 75.9 | 0.81% | 292/80 | 7.90% | 7.19 | 0.00318603 |
| 6 | Kola6 | LEM03 | modern | 201.5 | 41% | None | 10.67% | 197.0 | 0.86% | 380/86 | 10.80% | 17.95 | 0.00353702 |
| | Kola7 | LEM04 | modern | 206.3 | 41% | None | 9.98% | 201.8 | 0.88% | 381/86 | 11.70% | 18.06 | 0.00357139 |
| 7 | Kanin1 | SIW08 | modern | 177.9 | 41% | None | 11.24% | 172.2 | 1.33% | 375/92 | 15.10% | 12.47 | 0.00245719 |
| | Kanin2 | SIW09 | modern | 207.7 | 41% | None | 10.66% | 201.6 | 1.33% | 368/90 | 15.30% | 14.41 | 0.00231989 |
| 8 | NZemlya1 | 6350 | museum | 258.7 | 46% | None | 48.52% | 99.6 | 1.22% | 82/28 | 12.50% | 3 | NA |
| 9 | NZemlya2 | 6326 | museum | 50.6 | 44% | None | 32.06% | 48.5 | 1.13% | 109/41 | 7.60% | 1.97 | NA |
| 10 | NUrals | PS10k | ancient | 378.8 | 63% | 0,80% | 60.05% | 2.9 | 0.81% | 54/18 | 52.70% | 0.03 | NA |
| 11 | Ob' | 6357 | museum | 35.6 | 46% | 63% | 56.39% | 30.8 | 0.93% | 67/22 | 7.20% | 0.78 | NA |
| 12 | Yugorsky1 | 6359 | museum | 143.9 | 49% | 69% | 63.18% | 99.6 | 0.97% | 58/16 | 14.60% | 2.08 | NA |
| | Yugorsky2 | 6360 | museum | 20.3 | 49% | 49% | 66.65% | 13.8 | 0.88% | 53/14 | 8.80% | 0.27 | NA |
| 13 | Yamal1 | 662 | modern | 228.1 | 41% | None | 3.93% | 220.5 | 1.22% | 234/96 | 13.70% | 16.81 | 0.00641908 |
| 14 | Yamal2 | 391 | modern | 41.7 | 41% | None | 4.15% | 40.1 | 1.20% | 252/48 | 11.60% | 3.15 | 0.0050183 |

| | | | | | | | | | | | | | |
|---|---|---|---|---|---|---|---|---|---|---|---|---|---|
| | Yamal3 | 661 | modern | 80.1 | 40% | None | 5.24% | 77.5 | 1.24% | 301/77 | 8.30% | 7.42 | 0.00528077 |
| 15 | Gyda | 6328 | museum | 31.9 | 48% | 63% | 67.11% | 20.5 | 1.16% | 53/15 | 12.90% | 0.39 | NA |
| 16 | Yenisey | 6330 | museum | 207.9 | 45% | None | 37.94% | 205.1 | 1.20% | 98/38 | 12.60% | 7.44 | 0.01279665 |
| 17 | Taymyr1 | 6336 | museum | 112.2 | 39% | None | 3.61% | 93.4 | 1.30% | 215/58 | 11.30% | 6.98 | 0.00700459 |
| 18 | Taymyr2 | 6337 | museum | 220.0 | 45% | None | 29.27% | 99.4 | 1.30% | 119/42 | 11.20% | 4.38 | 0.01092966 |
| 19 | Taymyr3 | 6332 | museum | 203.3 | 46% | None | 37.09% | 200.7 | 1.35% | 99/36 | 12.70% | 7.33 | 0.01691918 |
| 20 | Lena | 369 | modern | 164.1 | 41% | None | 4.68% | 157.5 | 1.33% | 287/78 | 15.00% | 14.34 | 0.00749608 |
| 21 | Kotelny1 | 4991 | modern | 80.1 | 41% | None | 5.49% | 77.3 | 1.51% | 303/76 | 7.60% | 8.09 | 0.00527765 |
| | Kotelny2 | 4992 | modern | 80.0 | 41% | None | 5.06% | 77.3 | 1.49% | 297/73 | 6.80% | 7.39 | 0.00541258 |
| 22 | Verkhoyansk | 4464 | museum | 12.1 | 41% | None | 9.14% | 10.4 | 1.69% | 173/87 | 13.40% | 0.6 | NA |
| 23 | Indigirka1 | 6346 | ancient | 91.0 | 43% | None | 12.77% | 81.0 | 1.58% | 184/85 | 30.60% | 4.01 | 0.00704568 |
| | Indigirka2 | 6347 | ancient | 100.3 | 43% | None | 10.94% | 93.2 | 1.43% | 176/70 | 8.50% | 5.87 | 0.00711744 |
| 24 | Indigirka3 | 6380 | ancient | 406.3 | 51% | 2% | 18.78% | 7.6 | 1.58% | 104/46 | 6.20% | 0.3 | NA |
| 25 | Aldan | 4576 | museum | 315.9 | 43% | None | 4.84% | 98.1 | 1.36% | 143/60 | 24.20% | 5.02 | 0.00866139 |
| 26 | Zeya | 4568 | museum | 342.0 | 43% | None | 3.64% | 330.4 | 1.34% | 157/65 | 17.10% | 18.31 | 0.00519535 |
| 27 | Sikhote | 6320 | ancient | 201.7 | 62% | 2% | 15.04% | 3.9 | 1.32% | 118/45 | 53.90% | 0.07 | NA |
| 28 | Kamchatka1 | 396 | modern | 80.2 | 40% | None | 6.62% | 76.6 | 1.55% | 318/97 | 8.70% | 7.28 | 0.00289753 |
| 28 | Kamchatka2 | 397 | modern | 73.4 | 40% | None | 7.89% | 68.5 | 1.56% | 354/96 | 8.00% | 6.73 | 0.0027943 |
| 28 | Kamchatka3 | 398 | modern | 80.1 | 41% | None | 6.10% | 77.3 | 1.53% | 310/85 | 8.70% | 8.1 | 0.00316805 |
| 29 | Wrangel | SIE10 | modern | 232.9 | 41% | None | 9.17% | 224.8 | 1.53% | 360/89 | 13.00% | 19.09 | 0.00340783 |
| 30 | Magadan | 4541 | modern | 238.9 | 41% | None | 4.87% | 224.1 | 2.40% | 306/89 | 14.40% | 20.8 | 0.00652128 |
| 31 | Chukotka | 377 | modern | 206.8 | 41% | None | 4.04% | 195.5 | 2.46% | 269/86 | 18.20% | 16.55 | 0.00646922 |
| 32 | Alaska | TRI11 | modern | 221.0 | 41% | None | 9.11% | 208.5 | 2.50% | 354/95 | 13.00% | 17.27 | 0.00648969 |
| n/a | Dicr | 371 | modern | 84.4 | 42% | None | 4.07% | 65.7 | 8.58% | 267/75 | 14.30% | 4.99 | NA |

**Table S3. GADMA point estimates of demographic parameters for the best CLAIC model and the model with the highest log-likelihood, together with confidence intervals estimated for the best CLAIC model from bootstrap replicates.**

| Epoch | Parameters | best CLAIC model | Conf. intervals for the best CLAIC model | best logLL model |
|---|---|---|---|---|
| Ancestral | pop_size | 112125 | 106711 - 112242 | 110393 |
| Before split | pop_size end | 9256 | 4829 - 11670 | 20531 |
| | duration in ka | 1600 | 4830 - 11671 | 4708 |
| After split | pop_size start Kola | 5090 | 4247 - 7557 | 7048 |
| | pop_size end Kola | 71156 | 103425 - 408589 | 173398 |
| | pop_size start Scandinavia | 6855 | 41420 - 331228 | 109016 |
| | pop_size end Scandinavia | 166718 | 22546 - 56076 | 51740 |
| | duration in ka | 10748 | 8445 - 10693 | 5112 |
